# Knockout of the serotonin transporter in the rat mildly modulates decisional anhedonia

**DOI:** 10.1101/2020.07.08.190405

**Authors:** Chao Ciu-Gwok Guo, Michel MM Verheij, Judith R Homberg

**Affiliations:** Radboud University Medical Centre, Donders Institute for Brain, Cognition and Behaviour

**Keywords:** decisional anhedonia, cognitive effort, stimulus generalization, serotonin transporter knockout rat

## Abstract

Serotonin transporter gene variance has long been considered an essential factor contributing to depression. However, meta-analyses yielded inconsistent findings recently, asking for further understanding of the link between the gene and depression-related symptoms. One key feature of depression is anhedonia. While data exist on the effect of serotonin transporter gene knockout (5-HTT^-/-^) in rodents on consummatory and anticipatory anhedonia, with mixed outcomes, the effect on decisional anhedonia has not been investigated thus far. Here, we tested whether 5-HTT^-/-^ contributes to decisional anhedonia. To this end, we established a novel touchscreen-based “go/go” task of visual decision-making. During the learning of stimulus discrimination, 5-HTT^+/+^ rats performed more optimal decision-making compared to 5-HTT^-/-^ rats at the beginning, but this difference did not persist throughout the learning period. During stimulus generalization, the generalization curves were similar between both genotypes and did not alter as the learning progress. Interestingly, the response time in 5-HTT^+/+^ rats increased as the session increased in general, while 5-HTT^-/-^ rats tended to decrease. The response time difference might indicate that 5-HTT^-/-^ rats altered willingness to exert cognitive effort to the categorization of generalization stimuli. These results suggest that the effect of 5-HTT ablation on decisional anhedonia is mild and interacts with learning, explaining the discrepant findings on the link between 5-HTT gene and depression.

## Introduction

According to the World Health Organization (WHO), depression is the leading cause of disability worldwide, and a major contributor to the global burden of disease (Spencer L James, Degu Abate, Kalkidan Hassen Abate, Solomon M Abay, Cristiana Abbafati, et. al, 2018). More than 264 million people worldwide suffer from depression (Spencer L James, Degu Abate, Kalkidan Hassen Abate, Solomon M Abay, Cristiana Abbafati, et. al, 2018). Some people are more vulnerable to develop depression than others (Caspi et al., 2010), and multiple factors shape this vulnerability. The serotonin transporter (5-HTT) gene polymorphism presents one of these factors, specifically the short allelic variant of the serotonin-transporter-linked polymorphic region (5-HTTLPR). Some genetic meta-analyses studies reported statistical evidence for a relation between the 5-HTTLPR short allele and depression in the presence of life stress (Karg et al., 2011; Bleys et al., 2018) although others did not find this relationship (Culverhouse et al., 2018; Border et al., 2019). The discrepancies ask for a more behavioral approach and thus the identification of mechanisms that may link the 5-HTT gene to depression (Ormel et al., 2019). By regulating the function of the 5-HTT gene in animals, such as knocking out 5-HTT, it is possible to shed light on the nature of the potential relation between the 5-HTT gene and various aspects of depression.

Anhedonia is one of the essential features of depressive disorder and may bridge the link between the 5-HTT gene and depression. Anhedonia has been conceptualized in multiple aspects, including consummatory (“liking”), anticipatory (“wanting”), and decisional (“choosing”) anhedonia (Treadway and Zald, 2011; Der-Avakian and Markou, 2012; Ho and Sommers, 2013; Wu et al., 2017). While studies targeting anhedonia mostly address consummatory (“liking”) and anticipatory (“wanting”) anhedonia, decisional (“choosing”) anhedonia is less often considered. Decisional anhedonia of “choosing” highlights impaired decision-making in the context of reward and may refer to the impairment of balancing costs and benefits when selecting among multiple positive/rewarding options (Treadway and Zald, 2011). The role of serotonin in costs/benefits trade-off seems controversial: Pharmacological inhibition of 5-HTT reduced the expenditure of physical efforts in rats to maximizing rewards (Yohn et al., 2016) and blocking 5-HT synthesis had no effects (Denk et al., 2005). However, physical effort-based decision-making may differ from cognitive effort-based decision-making (Cocker et al., 2012). In addition, the options provided during decision-making are ambiguous in the real world (Lauriola and Levin, 2001; Huettel et al., 2006). Making decisions under ambiguity may trigger negative emotions, not only in humans (Rustichini, 2005) but also in animals (Nguyen et al., 2020). Whether the serotonin transporter plays an important role in cognitive effort-based decision-making remains to be determined, especially in animal studies (Silveira et al., 2020).

To test the relationship between the down-regulation of 5-HTT and anhedonia, 5-HTT^-/-^ rodents have been subjected to various anhedonia-related measurements. In an experiment where animals could freely choose in their homecage between water and a sucrose solution, 5-HTT knockout rats (Olivier et al., 2008), but not 5-HTT knockout mice (Kalueff et al., 2006) show increased consummatory (“liking”) anhedonia. In fixed ratio and progressive ratio tasks where rewards were presented to reinforce behavior, anticipatory (“wanting”) anhedonia-like behavior has been observed in 5-HTT^-/-^ mice (exert less physical effort to press the lever) (Sanders et al., 2007). Conversely, 5-HTT^-/-^ rats showed anticipatory addiction-like behavior when rewards were not presented to reinforce behavior (exert more physical efforts to press the lever) (Nonkes et al., 2010; Nonkes and Homberg, 2013). Taken together, the observations above suggest that 5-HTT^-/-^ rodents exert less physical efforts to acquire rewards if stimulus-reward associations remain stable, but exert more physical efforts to acquire rewards if stimulus-reward associations are devalued. These findings also indicate that 5-HTT deficiency in rats might affect the processing of reward in predicting conditioned stimuli. However, it is unclear whether and how 5-HTT deficiency affects the processing of the value properties of stimuli (e.g. subjective reward value) and/or the physical properties of stimuli (e.g. ambiguity). This insight is critical as anhedonia does not just involve hedonic capacity but also responsivity to pleasant stimuli (Der-Avakian and Markou, 2012).

In the current study, we designed a “reward/reward” paradigm of a “go/go” decision-making task in a touchscreen operant box, which maximizes the translational value into human studies. Both the responsivity to the reward value of stimuli and the ambiguity of the stimuli were precisely measured in this task. Rats were first required to discriminate between two reward conditioned stimuli (CSs, two different sizes of the white square): one CS was associated with a lower value of the reward, the other CS was associated with a higher value of the reward. To acquire the maximum amount of rewards, animals had to optimize their decision-making for each CS during stimulus discrimination. Animals suffering from decisional anhedonia may exert less cognitive effort to acquire the maximum amount of rewards. After reaching a criterion of discriminating between the two CSs correctly, rats were required to make decisions under the ambiguity of generalization stimuli (GSs). The relationship between CS and the GS, in the current task, is determined by the similarity of stimuli depending on the size of white squares. If the physical property (similarity or ambiguity) of a GS is close to one of the CSs, the decision on the GS is typically closer to the CS, resulting in a monotonic graded response to the stimuli (Ghirlanda and Enquist, 2003). The responsivity to the ambiguity of the GSs was measured during stimulus generalization. Generalizing the GSs to either lower reward value or higher reward value indicates the level of subjective value that the animals attribute to the stimulus. Rats suffering from decision anhedonia may attribute a lower subjective reward value to the GSs (Peters and Büchel, 2010). Similar “go/go” tasks have been established to test cognitive bias, which focused on negative versus positive interpretations of ambiguous information in rodents (Roelofs et al., 2016). These animal studies conceptualize both “reward/reward” and “reward/punishment” paradigms of the “go/go” decision-making in the same framework of cognitive bias (Roelofs et al., 2016; Nguyen et al., 2020). The information from the “reward/punishment” paradigm contains both negative and positive values and therefore the paradigm has been validated to test cognitive bias in depression as depressed subjects exhibit an information processing bias towards negative information (Gotlib et al., 2004; Joormann et al., 2006; Roelofs et al., 2016; Nguyen et al., 2020). However, anhedonia is not about the interpretation of negatively valenced stimuli, but rather about the interpretation of positively valenced stimuli. Therefore, to measure decisional anhedonia a “reward/reward” paradigm of the “go/go” decision-making task is most suited to test decision-making in the context of reward processing (Treadway and Zald, 2011; Der-Avakian and Markou, 2012).

We found that 5-HTT ablation may alter the responsivity to the reward value of stimuli during the learning of stimulus discrimination but may not alter the responsivity to the generalization stimuli. The results suggest that the effect of 5-HTT ablation on decisional anhedonia could be mild and interacts with the effect of learning. This mild effect could explain the discrepant findings on the link between the 5-HTT gene and depression. In sum, the absence of the 5-HTT might partially contribute to the development of decisional anhedonia.

## Materials and Methods

### Subjects

Ten wild-types (5-HTT^+/+^) and ten serotonin transporter knockout (5-HTT^-/-^) male Wistar rats aged between 68 and 84 days served as subjects. The total sample size of 20 rats in the current study was calculated based on visual reversal learning in 5-HTT^-/-^ versus 5-HTT^+/+^ rats (Nonkes and Homberg, 2013). Accordingly, we obtained a Partial η^2^ of 0.26191 (Lakens, 2013) and the effect size (0.55) was then determined. All rats were generated through homozygous breeding in our lab (Nijmegen, Netherlands). The homozygous parents were derived from heterozygous 5-HTT knockout rats that had been outcrossed for at least 10 generations with commercial Wistar Unilever rats (Charles River, Germany) (Homberg et al., 2007). All animals were housed in pairs under a reversed day/night cycle (light on from 19:00 to 7:00). From 7:00 to 19:00, animals were housed under dim red light conditions. The animals had *ad libitum* access to chow and water in their home cages. All experimental procedures were approved by the Central Committee on Animal Experiments (Centrale Commissie Dierproeven, CCD, The Hague, The Netherlands).

### Apparatus

Training and testing were performed in 10 operant chambers (Med Associates, USA) placed within sound-attenuated boxes. Each chamber was equipped with a pellet dispenser that delivered 45 mg sucrose pellet (TestDiet, USA) to a magazine when triggered. The boxes contained a magazine equipped with yellow light (ENV-200RL, Med Associates, USA) to indicate pellet delivery. A house light (ENV-215M, Med Associates, USA) for illuminating the chamber was mounted on the same wall above the magazine. The chambers contained a touchscreen positioned opposite the pellet dispenser. The screen was equipped with a metal plate containing three windows. See Fig. 1A for an illustration of a rat in the chamber facing the touchscreen presenting a stimulus image in the middle and choice images on both sides. The highest intensity of illumination in the operant chambers was about 35 lux (Lutron, LX-1108, TW). Computers were installed with K-limbic software which controlled the equipment and collected data from subjects’ responses.

**Fig. 1.**
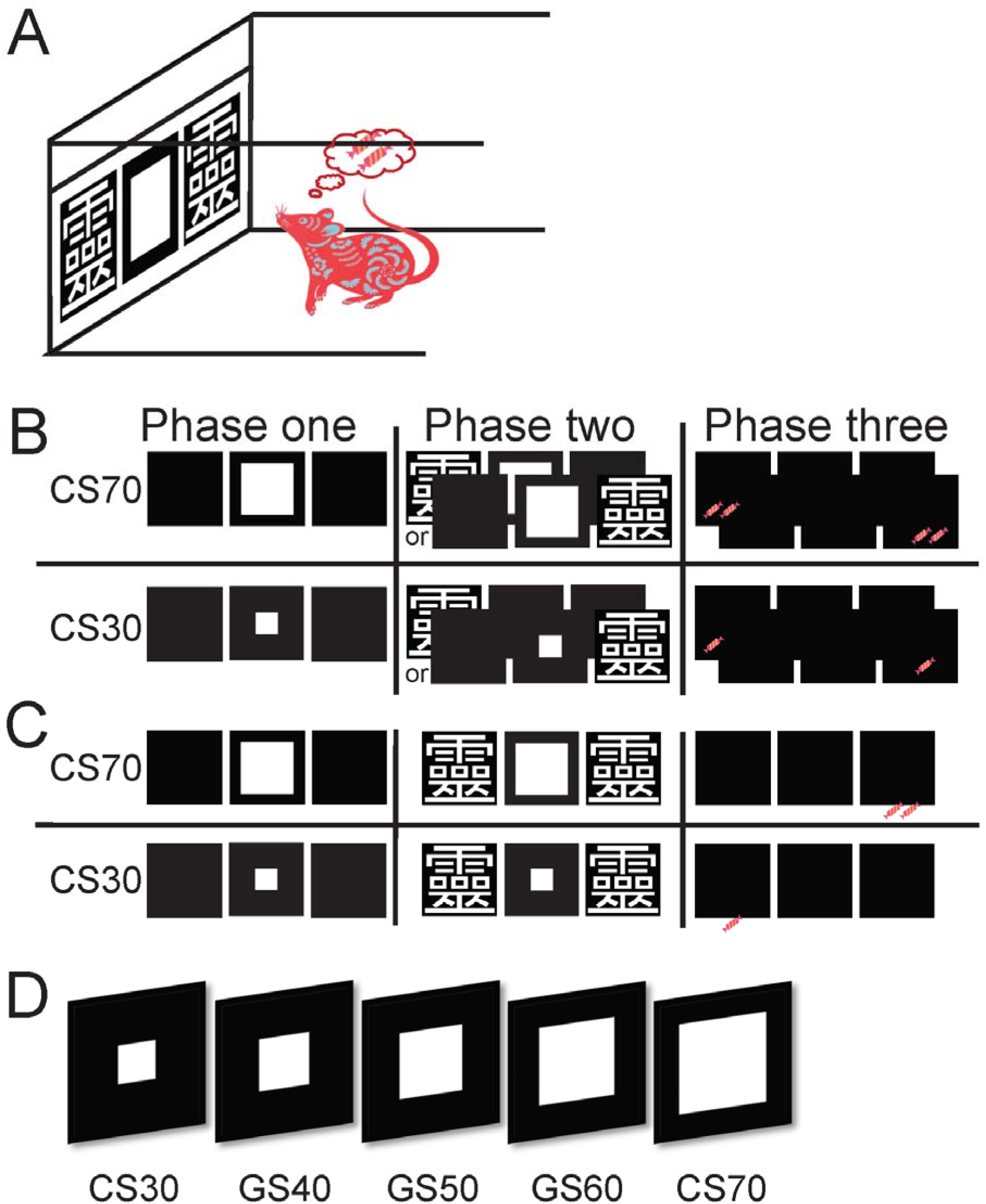
Schematic view of the experimental procedures of a rat. **(A)** A schematic view of a rat in an operant chamber with a touch screen. **(B) Sequential instrumental conditioning**. The sequence of events during CS30 and CS70 trials in the stage of instrumental conditioning. A rat was allowed to touch the white square image (phase one), which was followed by the random presentation of a reporter stimulus (RS) of 靈 on either the left or the right side of the screen (phase two). Then, a rat was allowed to touch this single RS, leading to the delivery of one sucrose pellet for CS30, and two sucrose pellets for CS70, stimuli (phase three). **(C) Stimulus discrimination**. The sequence of events during the CS30 and CS70 trials in the stage of stimulus discrimination. A rat was required to touch CS70 and a touch on the right-side RS only to acquire two sucrose pellets. The rat was required to touch CS30 and a touch on the left-side RS only to acquire one sucrose pellet. An incorrect response initiated a correction trial. **(D) Stimulus generalization**. A representation of all the stimuli during the stage of stimulus generalization. CS30 and CS70 are the same as above (see figures 1B and 1C). The white square image of the GS40 is larger than the CS30 while the white square image of the GS60 stimulus is smaller than that of the CS70. The size of GS50 is between the GS40 and GS60.

### Behavioral procedures

#### General

Each day before starting the experiment, rats stayed in a separate room for about 2 hours for habituation. At the same time, their chow was removed from their homecages. Rats had *ad libitum* access to chow in their homecages after behavioral training and testing. All rats were handled for 3 days before instrumental conditioning. The time to start testing was always between 13:00 and 14:00. Details about the conditioning are described as below:

#### Pre-training: Instrumental conditioning

Rats freely explored the touchscreen box during one session per day. Each session consisted of 30 trials. Each trial lasted for 30 seconds and trials were separated by a random inter-trial interval (ITI) between 15 and 30 seconds. During each trial, a small white circular disc (2.881*π* cm^2^) was presented on the screen in the middle of the central window. Each time the rat touched the circular disk, a sucrose pellet was dispensed into the magazine as a reinforcer. This trial was marked as a successful trial. Rats could constantly touch the disc and pellets were dispensed continuously within the period of a trial. If a rat did not touch the disc during the trial period, the houselight was illuminated for 5 seconds starting at the end of the trial as an indicator that the rat missed the trial. This was implemented to increase the attention of the rats to respond to stimuli. Rats received sequential instrumental conditioning (see below) once they completed at least 10 successful trials in the last session.

#### Pre-training: Sequential instrumental conditioning

Rats in this stage were trained to associate touching a conditioned stimulus (CS) and a reporter stimulus (RS) located at different positions of the screen sequentially with acquiring rewards. The CSs were either a white square of 25.2 or 10.8 cm^2^ (70% or 30% size of 36 cm^2^) and therefore were named CS70 and CS30 respectively. The RS was the Chinese character ‘靈’. Fig. 1B represents the trial sequence. Each trial contained three sequential phases. For example, CS70 was presented in phase 1, and if it was touched, RS was presented randomly on the left or right side in phase 2. The randomization of presenting the RS reduced the rats’ position bias to choose the left or right side of RS in the later stage of stimulus discrimination. If the RS was touched, two sucrose pellets were dispensed immediately in phase 3. Completion of a trial with a reward as an outcome was marked as a successful trial. If the trial was omitted, the next trial was presented. Nothing happened if a rat touched the wrong screen before or after the RS was presented. CS30 trials were similar to CS70 trials, except that rats were rewarded with one sucrose pellet in each successful trial. Each trial lasted for 40 seconds and trials were separated by a random range of ITIs lasting between 15 seconds and 30 seconds. If a rat omitted a trial (the CS70 or the RS was not touched during the trial period), the houselight was illuminated for 5 seconds at the end of the trial. During ITI, nothing happened if a rat touched the screen. Rats were subjected to 20 CS70 trials and 20 trials CS30 per session in total and received one session per day. The order of each trial type was randomly assigned. CSs associated with one or two sucrose pellets were counterbalanced across groups so that half of the CS70 trials were associated with two sucrose pellets and the other half of CS30 trials were associated with one sucrose pellet. After receiving five consecutive sessions, rats were shifted to the next stage of discrimination conditioning.

#### Stimulus discrimination

In this stage, rats had to exert efforts to memorize the discrepancy between CS70 and CS30 to acquire the rewards. To earn rewards, they had to make decisions based on the CS temporarily presented within a trial. Each trial contained three sequential phases. Fig. 1C represents a trial sequence. As an example, when a rat was tested during a CS70 trial, the CS70 was presented in phase 1, and if it was touched, two RSs were presented on both left and right sides in phase 2. Subsequently, if the RS was presented on the right side of the screen was touched, two sucrose pellets were immediately dispensed in phase 3. Completing a trial by acquiring a reward was marked as a successful trial. The sequence of phases for the CS30 trial was similar as for the CS70 trial, except that rats were rewarded with one sucrose pellet in a successful trial by touching the RS presented on the left side of the screen (see Fig 1C, CS30). Correction trials were applied if rats touched the wrong RS or omitted the trial. During correction trials, the same trial was repeated until rats made the correct response. Both the CS and the position of the target RS were counterbalanced across the rats, such that for half of the rats the RS presented on the left side was the target stimulus and the RS on the right side was the dummy stimulus during CS30 trials, while the target and dummy RSs were opposite during CS70 trials. For the other half rats, the RS presented on the right side was the target stimulus during CS30 trials and on the left side during CS70 trials. Rats were subjected to 40 trials (20 CS30 and 20 CS70 trials) per session. The order of the two CS trials was randomized in each session and no more than four consecutive trials were the same. Rats received one session per day until they correctly completed at least 70% of trials in each session of the last three consecutive sessions.

#### Stimulus generalization

The animals received three identical generalization sessions, conducted on three separate days. To verify that the discrimination accuracy had not declined before each generalization test, the discrimination accuracy of the CSs was tested between the generalization tests. If accuracy was less than 70%, rats were exposed to more sessions of stimulus discrimination until the accuracy was higher or equal to 70% at the last two sessions before the next day of the generalization test. In the stimulus generalization test, two CSs and three GSs were introduced. The square size of the GSs was 40% (GS40), 50% (GS50), and 60% (GS60) of 36 cm^2^, respectively. The sequence of GS trials was similar to CS trials except that no reward outcome was delivered in phase 3 to reduce a reward conditioned effect on the ambiguity of the GSs. Fig.1 D presented the CS and GS trials applied during the stimulus generalization test. Each session consisted of 72 trials, in which GS40, GS50, GS60 each was presented during 12 trials and CS30, CS70 each during 18 trials. The 72 trials were divided into 12 blocks resulting in 6 trials per block. The inter-block interval was one minute. The order of the trials in each block was a CS followed by a GS to guard against the sequential effects of the same stimulus. The CSs and GSs were evenly distributed in each block and counterbalanced across blocks and groups.

### Data analysis

The number of sessions needed to reach the discrimination criterion was statistically analyzed using a t-test with python in DABEST (version 0.3.1). The genotype effect distribution was estimated via 5000 times of bootstrapping (Ho et al., 2019). The rest of the data were analyzed using a mixed-effect logistic or linear regression model by software R (version 3.6.2) with package lme4 (Bates et al., 2015). The mixed-effect modeling analysis reduces the probability of false positives (Aarts et al., 2014). Genotype, stimuli, and session were entered as fixed effects, and subjects and the date of the experiment were entered as random effects in the model. The linear regression model equation predicts the outcome (*Y*) as follows:

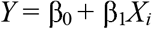

where β_0_ is the intercept and β1 is the slope of the independent variable X.

The logistic regression model equation predicts the odds ratio of a successful event (reached 70% accuracy in a session) as follows:

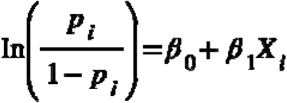

where *p* is the probability of a successful event, which indicates the proportion of successful events (reached 70% accuracy) at any given number of the session.

Visual inspection of the residuals distribution and residual Quantiles-Quantiles (Q-Q plot) were used to check the heteroskedasticity of data. Q-Q plot is a graphical technique for determining if the data we collected come from populations with a normal distribution. If the data set comes from a normal distribution, the data points fall approximately along a diagonal solid line. The estimates of the fixed effect distributions were obtained by bootstrapping 5000 times and plotted with R (version 3.6.3). The 95% of bootstrapped confidence intervals (BCI) and the estimates were provided for statistical inference. If the bootstrapped 95% CI did not include 0, the estimated effect was designated to be “significant”. Also, we described the effect as “strong”, “moderate” or “mild” level based on the distribution via visual inspection. All the intercept effects from each model are “strong”. Individual data points and regression curves were plotted with python (version 3.7).

## Results

### Stimulus discrimination

Rats had to exert efforts to memorize the discrepancy between CS30 and CS70 to acquire the maximum number of rewards by making optimal decisions when a single CS was temporally presented. Their decisions resulted in a discrimination performance during a session, which we termed the discrimination accuracy. It is defined as the percentage of correct trials in a session (correction trials were not included). The performance of discrimination accuracy across the sessions is presented in Fig.2 A1. A t-test indicated that 5-HTT^-/-^ and 5-HTT^+/+^ rats needed a similar amount of sessions to reach the criterion of 70% accuracy of pooled CSs for consecutive three sessions (E = -2.3, BCI = [-5.7, 1.5], see Fig.2 A2). Although the accuracy for pooled CSs across 3 consecutive sessions was 70%, rats achieved an accuracy higher in CS70 than in the CS30 trials (see Fig.2 B1), and the stimulus effect was “significant” (E = -11.2083, BCI = [-12.83, - 9.52], Fig.2 B2). The intercept of the model was also “significant” (E = 77.2917, BCI = [75.65, 78.95], Fig.2 B2). There were no other “significant” effects (genotype: E = -0.3750, BCI = [2.0059, 1.2936]; session: E = 0.4375, BCI = [-1.5887, 2.4885]; genotype*session: E = 0.1875, BCI = [-1.8103, 2.2323]; see Fig.2 B2). The residuals of the data were normally distributed (residuals distribution, Fig.2 C1; Q-Q distribution, Fig.2 C2). CSs were associated with either one sucrose pellet or two sucrose pellets. To acquire the maximum number of pellets, rats had to make optimal decisions across sessions during both CSs trials. Logistic regression was performed to assess rats’ performance of discrimination accuracy for each of the CS30 and CS70 trials. The mixed-effect logistic regression model showed that session, session*genotype, stimulus, intercept effects were “significant”, and that genotype and genotype*stimulus effect were not “significant” (genotype: -0.07378 BCI = [-0.6298, 0.4668]; genotype*stimulus: E = -0.15606, BCI = [-0.4654, 0.1654]; session: E = 0.31154, BCI = [0.2302, 0.4233]; session*genotype: E = 0.07383, BCI = [0.0052, 0.1567]; stimulus: E = 1.63932, BCI = [1.355, 2.033]; intercept: E = - 0.61645, BCI = [-1.2811, -0.0101], Fig.3 A). The residuals of the data seem normally distributed (Fig.3 B1) and most data residuals are within the 95% of the theoretical normal distribution (Fig.3 B2). For the CS trials associated with higher reward value (CS70) (see Fig.3 C1), the intercept and session effects were “significant” and the genotype and genotype*session effects were not “significant” (genotype: E = -0.32324, BCI = [-1.2618, 0.5338]; session: E = 0.59286, BCI = [0.4602, 0.8623]; genotype*session: E = 0.03142, BCI = [-0.1345, 0.2112]; intercept: E = 1.60289, BCI = [0.845, 2.691]; Fig.3 C2). For the CS trials associated with one sucrose pellet (CS30) (Fig.3 D1), intercept and session effects were “significant” and the genotype and genotype*session effects were not “significant” (genotype: E = -0.01332, BCI = [-0.7609, 0.7191]; session: E = 0.18242, BCI = [0.0819, 0.3087]; genotype*session: E = 0.06724, BCI = [-0.0289, 0.1792]; intercept: E = -2.02776, BCI = [3.067, -1.325], Fig.3 D2).

**Fig. 2.**
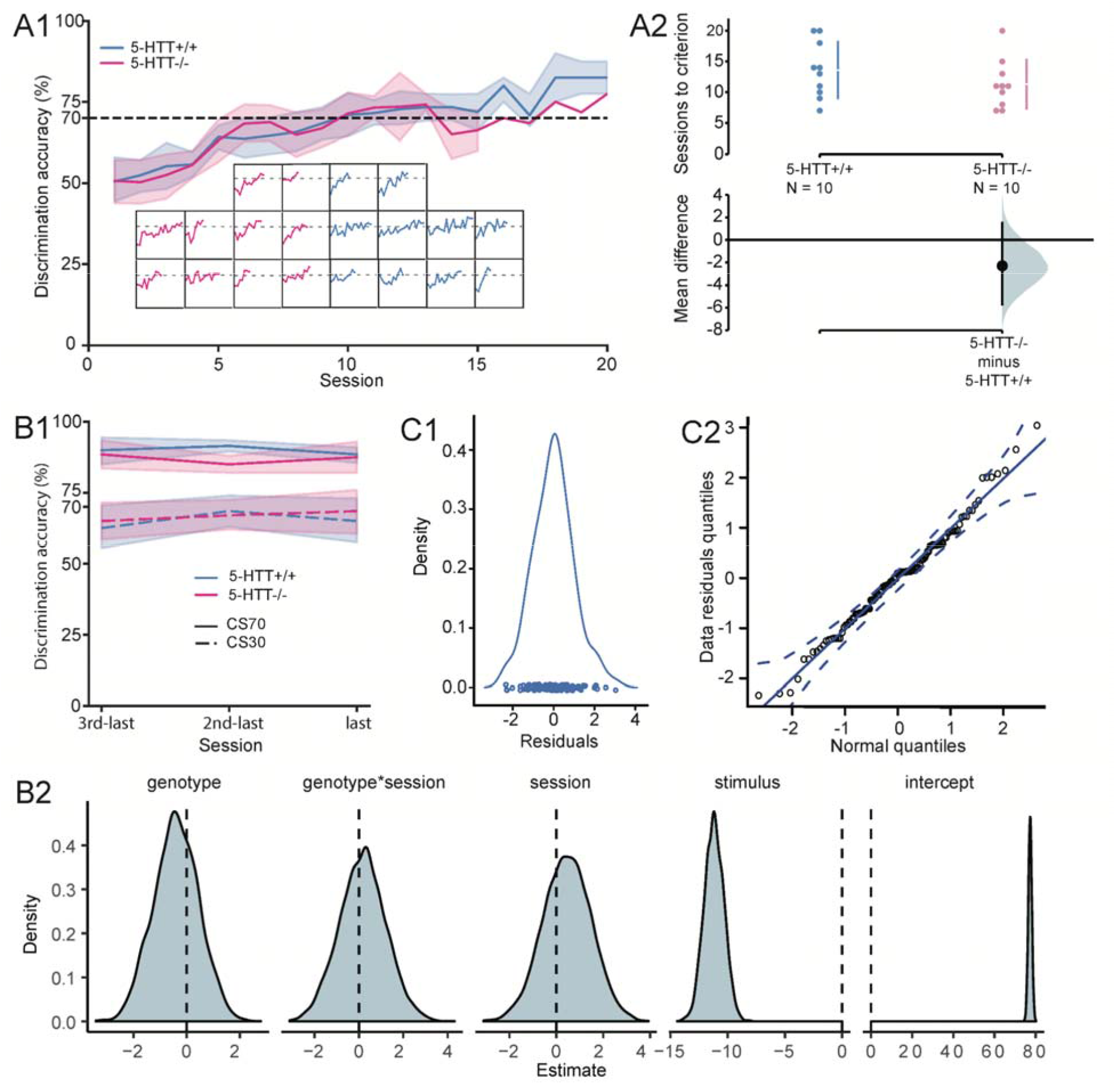
The accuracy of stimulus discrimination. **(A1)** Solid lines represent the mean of discrimination accuracy of pooled CSs in each genotype. The transparent stripes are 95% confidence intervals. Figures embedded in panel A are the individual performance data for discrimination accuracy of pooled CSs in every session. **(A2)** The number of sessions the rats needed to reach the learning criterion. The effect size and the estimation of 95% confidence intervals obtained from bootstrapping are plotted on separate axes beneath the individual data points. For each genotype, mean⍰± ⍰ standard deviations are shown as vertical gapped lines. **(B1-B2)** The accuracy of the last three sessions during stimulus discrimination. **(B1)** Blue and red solid curves represent the mean accuracy for CS70 and blue and red dashed curves the mean accuracy for CS30. The transparent strips around the mean are 95% confidence intervals. **(B2)** The estimation of each fixed effect by bootstrapping 5000 times from the mixed-effect linear model. The grey shaded area is the probability of the estimation of the effect. The stimulus effect is “strong”. **(C1-C2)** Data distribution. **(C1)** The distribution of residuals of the response accuracy during the last three sessions of stimulus discrimination. A circle represents a residual of a data point. The distribution of the data points follows a normal assumption. **(C2)** The quantile-quantile distribution. The diagonal solid line represents a normal distribution. A circle represents an observed data point against a data point from a normal distribution. The dashed lines present the lower limit and upper limit of the 95% confidence interval. The data points fall approximately along the solid line.

**Fig. 3.**
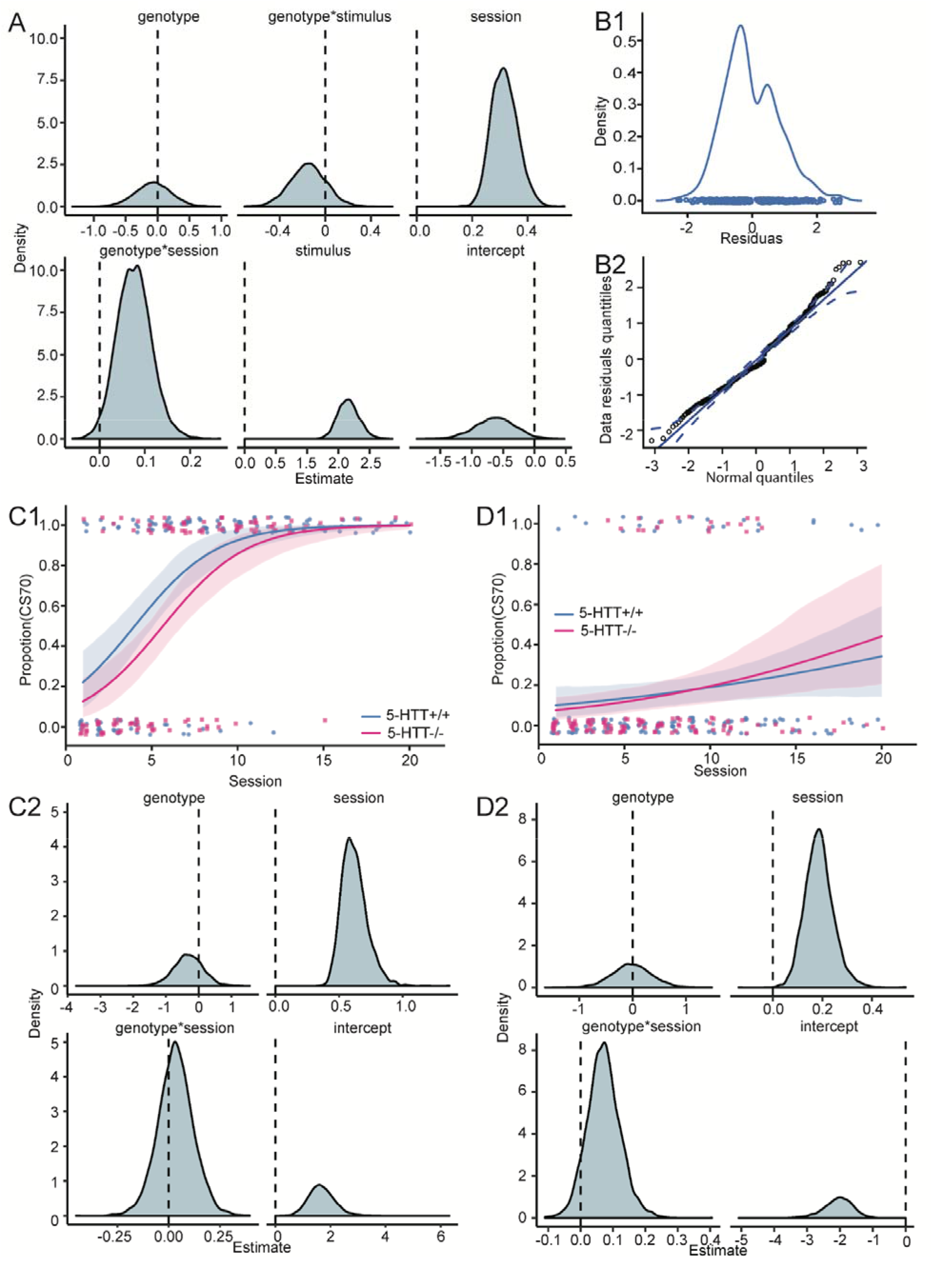
The proportion of reaching the 70% accuracy during the learning of stimulus discrimination. **(A-B)** Stimulus discrimination for both CS30 and CS70. **(A)** The estimation of each fixed effect by bootstrapping 5000 times from the mixed-effect logistic model. The grey shaded area is the probability of the estimation of the effect., genotype*session effect is “moderate”. Session and stimulus effects are “strong”. **(B1-B2)** Data distribution. **(B1)** The distribution of residuals of the proportion of successful events during stimulus discrimination. A circle represents a residual of a data point. The distribution of the data points approximately follows a normal assumption. **(B2)** The quantile-quantile distribution. The diagonal solid line represents a normal distribution. A circle represents an observed data point against a data point from a normal distribution. The dashed lines present the lower limit and upper limit of the 95% confidence interval. The data points fall approximately along the solid line. **(C1-C2)** Stimulus discrimination of CS70. (C1) Logistic regression curves for CS70. The solid lines are the logistic regression lines. A dot in the top denotes a subject that reached 70% accuracy in a session, a dot in the bottom denotes a subject that failed to reach 70% accuracy in a session. The transparent stripes denote the 95% confidence intervals of the regression lines. **(C2)** The estimation of each fixed effect by bootstrapping 5000 times from the mixed-effect logistic regression model. The grey shaded area is the probability of the estimation of the effect. The session effect is “strong”. **(D1-D2)** Stimulus discrimination for CS30. **(D1)** Logistic regression curves for CS30. The solid lines are regression lines. A dot in the top denotes a subject that reached 70% accuracy in a session, a dot in the bottom denotes a subject that failed to reach 70% accuracy in a session. The transparent stripes denote the 95% confidence intervals of the regression lines. **(D2)** The estimation of each fixed effect by bootstrapping 5000 times from the mixed-effect logistic model. The grey shaded area is the probability of the estimation of the effect. The session effect is “strong” and the genotype*session effect is “mild”.

In summary, both 5-HTT^+/+^ and 5-HTT^-/-^ rats had a higher accuracy for CS70 than CS30, and stimulus effects (Fig.2 B and Fig.3 A) were “significant”. The interaction effect of the genotype*session was “significant” in the full mixed-effect logistic model (Fig.3 A), but the effect in the logistic model for the CS70 and CS30 stimuli separately was not “significant” (Fig. 3 C2 and D2). These results indicate that the difference between 5-HTT^+/+^ and 5-HTT^-/-^ rats for more optimal decision-making was mild and that 5-HTT^+/+^ rats might perform more optimal decision-making at the beginning of discrimination, but this effect did not persist. Importantly, both genotypes increased the discrimination accuracy as the session increased (Fig.3 A, C, D) and were able to reach the 70% accuracy criterion of pooled CSs across three consecutive sessions similarly (Fig. 2 A).

### Stimulus generalization

Rats generalize the learned information from CS30 and CS70 to GS40, GS50, and GS60 based on the physical property (similarity or ambiguity) among the stimuli by making decisions on the stimulus currently presented on the screen. If the physical property of a GS is close to one of the CSs, the decision on the GS is typically closer to the CS, resulting in a monotonic graded response to the stimuli (Ghirlanda and Enquist, 2003). The mixed-effect linear model shows that there was a “significant” effect of stimulus similarity on decision-making (E = 15.0370, BCI = [13.91, 16.17], Fig.4 A). The intercept effect of the model is also “significant” (E = 55.7362, BCI = [51.66, 59.80], Fig.4 A). However, genotype, genotype*session, genotype*stimulus, session effects were not “significant” (genotype: E = 1.1945, BCI = [-1.863, 4.343]; genotype*session: E = 0.5345, BCI = [-1.6028, 2.6864]; genotype*stimulus: E = -0.5741, BCI = [-1.7058, 0.5737], session: E = 0.2927, BCI = [-2.3067, 2.8971]). The residuals of data are normally distributed (Fig.4 B1 and B2). As the physical property (similarity) was closer to CS70, the decisions were more robust in general and in each session (the percentage of categorizing the stimulus as CS70 as the target, see Fig.4 A, C1). However, the percentage of responses to CS70 (the percentage of categorizing the stimulus as CS70) was similar between sessions in general and in each stimulus (See Fig.4 A, C2). The individual response (percentage of responses to CS70) of stimulus generalization in each session is shown in Fig.4 C3. We observed six types of stimulus generalization curves: monotonic grade (orange background), ‘S’ shape (green background), monotonic peak at GS60 (banana background), monotonic peak at GS40 (brown background), GS50 as a peak (coral background) and GS50 as a peak valley (charcoal background). The yellow background of the figure indicates a monotonic graded response. The major type of generalization curve is monotonic graded responses, which was found for 17/30 sessions in 5-HTT+/+ rats, and 14/30 sessions in 5-HTT-/- rats. Regarding the number of rats, 3/10 5-HTT^-/-^ rats had at least two sessions of monotonic graded responses, while 5/10 5-HTT^+/+^ rats had at least two sessions of monotonic graded responses. Notably, the percentage of response to CS70 is higher than CS30 of each rat at each session.

**Fig. 4.**
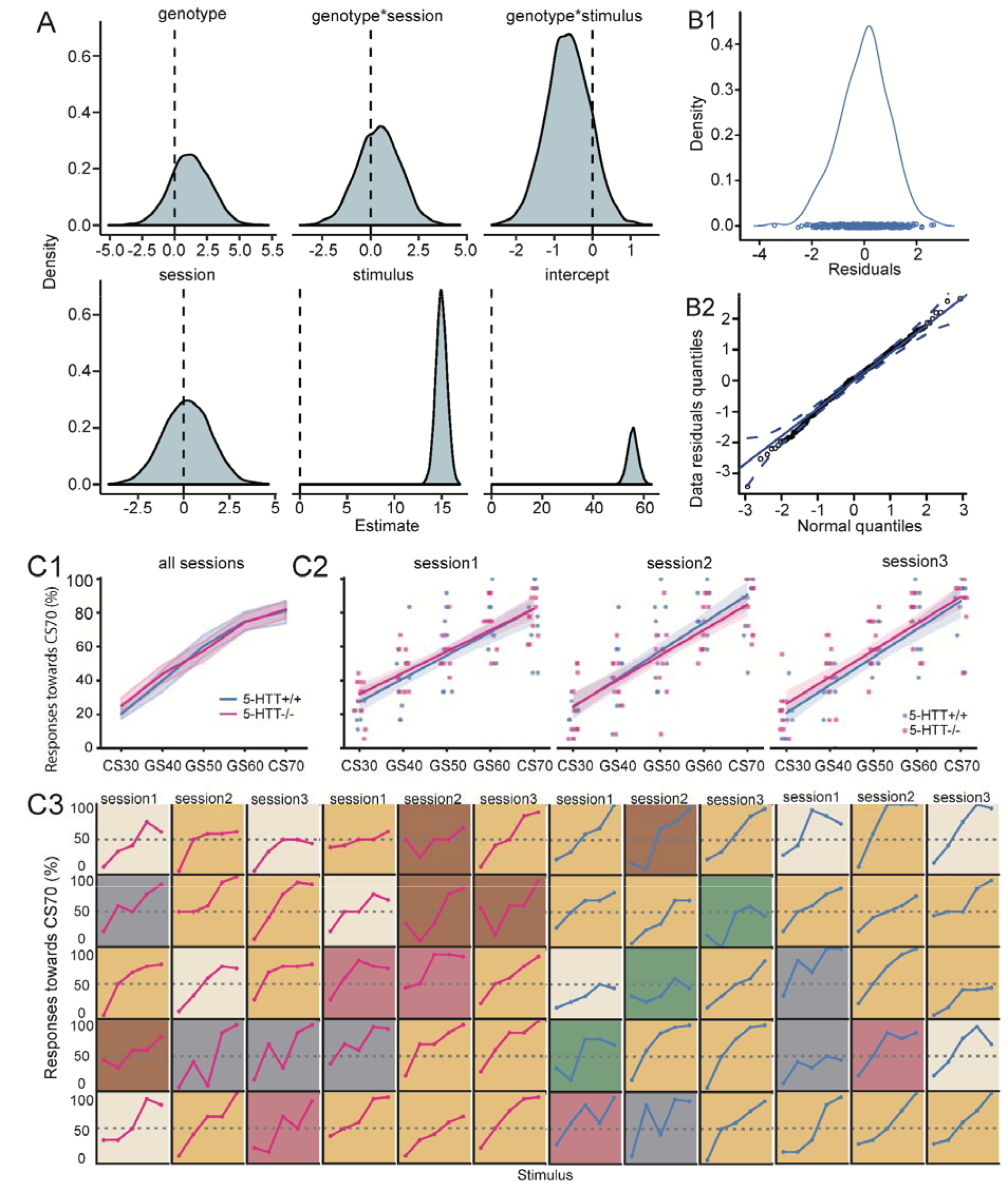
Stimulus generalization. The decisional anhedonia of responding to GSs was similar between 5-HTT^-/-^ and 5-HTT^+/+^ rats. **(A)** The estimation of each fixed effect by bootstrapping 5000 times from the mixed-effect linear model. The grey shaded area is the probability of the estimation of the effect. The stimulus effect is “strong”. **(B1-B2)** Data distribution. **(B1)** The distribution of residuals of the percentage of responses towards CS70. A circle represents a residual of a data point. The distribution of the data points follows a normal assumption. **(B2)** The quantile-quantile distribution. The diagonal solid line represents a normal distribution. A circle represents an observed data point against a data point from a normal distribution. The dashed lines present the lower limit and upper limit of the 95% confidence interval. The data points fall approximately along the solid line. (C1-C3) Stimulus generalization curves **(C1)** The percentage of responses towards CS70 increased with the similarity of the GS closer to the CS70. Solid lines represent the mean percentage of choices towards CS70. Stripes represent 95% confidence intervals of the mean. **(C2)** The percentage of responses towards CS70 in each session. The solid lines are linear regression lines. The transparent stripes denote the 95% confidence intervals of the regression lines. A dot represents a percentage of trials in a session that categorizing the stimulus as CS70. **(C3) Stimulus generalization in individuals**. A single line represents a generalization curve of a rat per session. Red lines represent the generalization curve of 5-HTT^-/-^ rats, blue lines represent the response curve of 5-HTT^+/+^ rats. Six types of stimulus generalization curves are denoted by the colour of the background: monotonic grade (orange background), ‘S’ shape (green background), monotonic peak at GS60 (banana background), monotonic peak at GS40 (brown background), GS50 as a peak (coral background) and GS50 as a peak valley (charcoal background).

Before taking optimal actions on the stimulus, the response time (RT) plays a crucial role in information processing both in rats and humans (Donders, 1969; Blokland, 1998). We, therefore, further analyzed rats’ response time to the stimuli. A mixed-effect linear model showed that genotype, genotype*stimulus, session effects were not “significant” (genotype: E = 0.07179, BCI = [-0.3388, 0.4630]; genotype*stimulus: E = -0.0004043, BCI = [-0.1396, 0.1322]; session: E = 0.09016, BCI = [-0.2202, 0.3834]; Fig.5 A). Genotype*session, stimulus and intercept effects were “significant” on the RT (genotype*session: E = -0.40600, BCI = [-0.6641, -0.1492]; stimulus: E = 0.46803, BCI = [0.3355, 0.6052]; intercept: E = 4.10327, BCI = [3.608, 4.606]; Fig.5 A, C and D). Since genotype*session effect was “significant”. A follow-up analysis of mixed-effect linear models for each genotype was performed. For 5-HTT^+/+^ rats, the intercept effect, stimulus and session effects were “significant” (intercept: E = 4.0025, BCI = [3.336, 4.689]; stimulus: E = 0.4684, BCI = [0.3140, 0.6196]; session: E = 0.5320, BCI = [0.1878, 0.8883]; Fig.5 D2). For 5-HTT^-/-^ rats, the intercept and stimulus effects were “significant” (intercept: E = 4.2197, BCI = [3.500, 4.901]; stimulus: E = 0.4676, BCI = [0.2596, 0.6800]; Fig.5 D2), while session effect was not significant (E = -0.4199, BCI = [-0.9242, 0.0813]; Fig.5 D2).

**Fig. 5.**
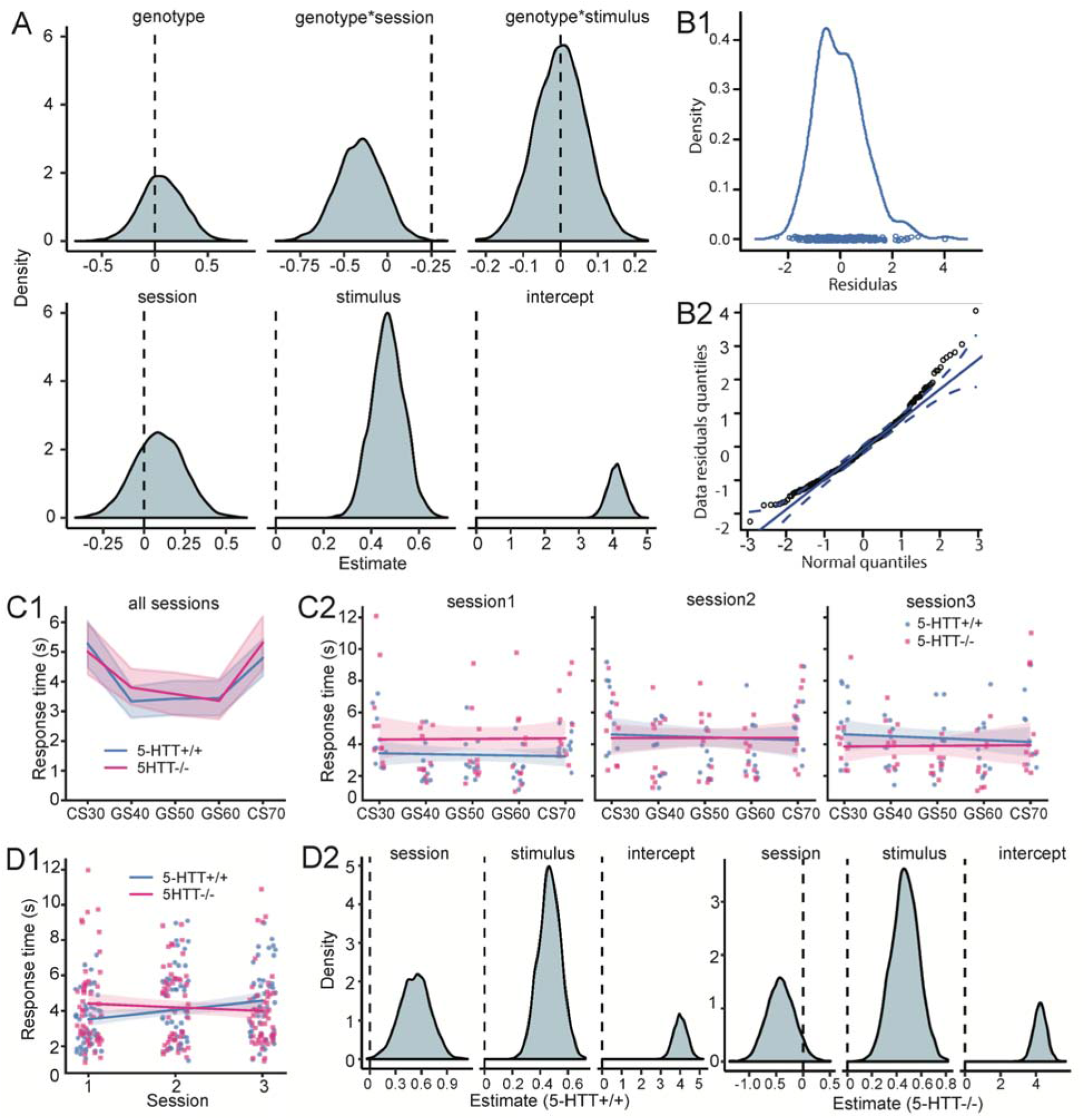
Response time (RT) during the stimulus generalization. Response time changes differently between 5-HTT^+/+^ and 5HTT^-/-^ rats. **(A)** The estimation of each fixed effect by bootstrapping 5000 times from the mixed-effect linear model. The grey shaded area is the probability of the estimation of the effect. Genotype* session and stimulus effects are “strong”. **(B1-B2**) Data distribution. **(B1)** The distribution of residuals of the RT. A circle represents a residual of a data point. The distribution of the data points approximately follows a normal assumption. **(B2)** The quantile-quantile distribution. The diagonal solid line represents a normal distribution. A circle represents an observed data point against a data point from a normal distribution. The dashed lines present the lower limit and upper limit of the 95% confidence interval. The data points fall approximately along the solid line. **(C1-C2)** RT for each stimulus. **(C1)** RT for each stimulus in all sessions. Solid lines represent the mean of RT. Stripes are 95% confidence intervals around the mean of RT. **(C2)** RT for each stimulus in each session. The solid lines are linear regression lines. The transparent stripes denote the 95% confidence intervals of the regression lines. A dot represents an RT of a rat to a stimulus. **(D1-D2)** RT of all stimuli in each session. **(D1)** The solid lines are the linear regression with the transparent stripes of 95% confidence intervals around. A dot denotes the individual RT in a session. **(D2)** The estimation of each fixed effect by bootstrapping 5000 times from the mixed-effect linear model in 5-HTT^-/-^ rats and 5-HTT^+/+^ rats separately. The grey shaded area is the probability of the estimation of the effect. Session and stimulus effects are “strong” in 5-HTT^+/+^ rats. In 5-HTT^-/-^ rats, the session effect is “mild” and the stimulus effect is “strong”.

Since each stimulus presented to the rats was different, mixed-effect linear models for each stimulus were then performed. For CS30 (Fig.6 A1), the intercept effect was “significant” (intercept: E = 5.1449, BCI = [4.572, 5.705]; Fig.6 A2); while genotype and session effects were not “significant” (genotype: E = -0.13505, BCI = [-0.7042, 0.4420]; genotype*session: E = - 0.56541, BCI = [-1.2578, 0.1038]; session: E = 0.05031, BCI = [-0.6316, 0.7381]; Fig.6 A2). Notably, Fig.6 A2 shows that the genotype*session might have a mild “significant” effect on RT. For GS40 (Fig.6 B1), session and intercept effects were “significant”, and the other effects were not “significant” (genotype: E = 0.23112, BCI = [-0.2298, 0.7168]; genotype*session: E = - 0.06797, BCI = [-0.5248, 0.3938]; session: E = 0.45555, BCI = [-0.0106, 0.9236]; intercept: E = 3.56528, BCI = [3.091, 4.029]; Fig.6 B2). For GS50 (Fig.6 C1), the intercept effect was “significant”, and the other effects were not “significant” (genotype: E = 0.07775, BCI = [-0.4385, 0.5782]; genotype*session: E = -0.43757, BCI = [-0.9395, 0.0632]; session: E = 0.02282, BCI = [-0.4751, 0.5325]; intercept: E = 3.50214, BCI = [2.995, 4.034]; Fig.6 C2). Notably, Fig.6 C2 shows that the genotype*session might have a mild “significant” effect on RT. For GS60 (Fig.6 D1), genotype*session and intercept effects were “significant”, and the other effects were not “significant” (genotype: E = -0.04251, BCI = [-0.6394, 0.5184]; genotype*session: E = -0.56101, BCI = [-1.0383, -0.0849]; session: E = 0.10547, BCI = [-0.3578, 0.5993]; intercept: E = 3.39427, BCI = [2.839, 3.981]; Fig.6 D2). For CS70 (Fig.6 E1), intercept effect was “significant” and the other effects were not “significant” (genotype: E = 0.2588, BCI = [-0.4163, 0.9159]; genotype*session: E = -0.2262, BCI = [-0.8683, 0.3961]; session: E = 0.1374, BCI = [-0.4791, 0.7422]; intercept: E = 5.0525, BCI = [4.410, 5.706]; Fig.6 A2).

**Fig. 6.**
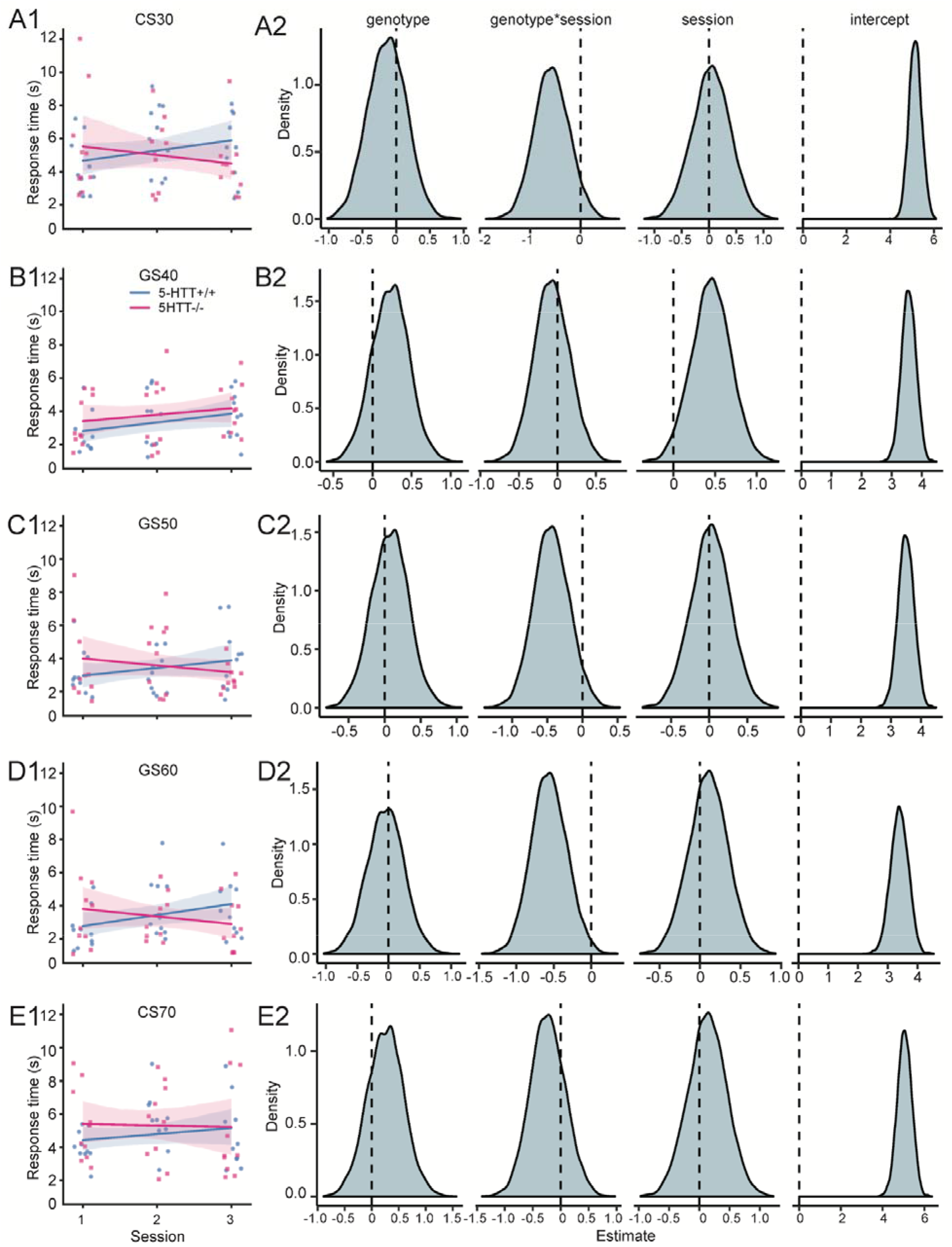
The RT of each stimulus during stimulus generalization. **(A1-A2)** RT of CS30 in each session. The genotype*session effect is “mild”. **(B1-B2)** RT of GS40 in each session. Session effect is “mild”. **(C1-C2)** RT of GS50 in each session. The genotype*session effect is “mild”. **(D1-D2)** RT of GS60 in each session. The genotype*session effect is “moderate”. **(E1-E2)** RT of CS70 in each session. Note: The solid lines are the linear regression lines, surrounded by transparent stripes of 95% confidence intervals. A dot denotes the individual response time in a session. The distribution is the estimation of each fixed effect by bootstrapping 5000 times from the mixed-effect linear model. The grey shaded area is the probability of the estimation of the effect.

Taken together, 5-HTT^+/+^ rats tend to increase RT across the three generalization test sessions while 5-HTT^-/-^ rats tend to decrease RT. However, the generalization curves were similar between the genotypes.

## Discussion

We found that both 5-HTT^-/-^ and 5-HTT^+/+^ rats were able to reach 70% accuracy across three consecutive sessions during stimulus discrimination. Also, the number of sessions needed to reach such a criterion was similar between the two genotypes. However, 5-HTT^+/+^ rats performed more optimal decision-making compared to 5-HTT^-/-^ rats during stimulus discrimination at the beginning, but this effect didn’t persist as the session progressed. There was an interaction between genotype and learning sessions. Further, we found that 5-HTT^+/+^ rats gradually increased response time to generalization stimuli across the three generalization test sessions, while 5-HTT^-/-^ rats only tended to decrease response time. The stimulus generalization curves were similar between the two genotypes and did not alter as the sessions progressed.

During stimulus discrimination, all 5-HTT^-/-^ and 5-HTT^+/+^ rats reached 70% accuracy across the last three consecutive sessions, which indicates that both genotypes were able to exert efforts to discriminate between the two CSs with similar accuracy. Interestingly, rats performed better in CS70 (higher reward) than CS30 (lower reward) trials. We assumed that rats could perform similar accuracy in both CS70 and CS30 during the learning of discrimination. The discrepant accuracy between CS30 and CS70 indicates that rats overall did not maximize their cognitive efforts to discriminate between stimuli for the maximum amount of rewards. Similarly, humans also overall tend to avoid exerting cognitive efforts (Otto and Daw, 2019). Whether this discrepancy of lower and higher rewards can be eliminated through continuous training needs further research. In other similar experiments, different accuracy criteria were applied (for example, accuracy for each of the two CSs is higher than 60% or accuracy for two CSs pooled is higher than 80%) (Hales et al., 2016; Krakenberg et al., 2019a, 2019b). Even when the higher criterion was set at 80% accuracy for stimulus discrimination, no genotype differences were found between 5-HTT^-/-^ and 5-HTT^+/+^ rodents (Krakenberg et al., 2019a). However, whether there was a genotype difference during the course of stimulus discrimination was not reported in mice lacking 5-HTT^-/-^ (Krakenberg et al., 2019a).

During the early sessions of stimulus discrimination, more 5-HTT^-/-^ rats displayed lower learning accuracy than 5-HTT^+/+^ rats. As the session increased, both 5-HTT^-/-^ and 5-HTT^+/+^ rats reached similar discrimination accuracy at the end. This indicates that 5-HTT^-/-^ rats have a reduced capability to balance the cost of efforts and benefit of reward outcome during stimulus discrimination (i.e. decisional anhedonia) in the early phase. This interaction effect, however, was not observed when we analyzed the responses to CS30 and CS70 separately. In our current study, 5-HTT^-/-^ rats, compared with 5-HTT^+/+^ rats, might exert less cognitive effort to acquire reward in the early phase of stimulus discrimination, but exert more cognitive effort to acquire reward in the later phase. Fewer studies are focusing on the function of serotonin in cognitive efforts. One pharmacological study in rats published recently found that inhibiting or activating 5-HT1A, 5-HT2A, and 5-HT2C receptor subtypes didn’t shift rats’ willingness to exert cognitive effort for larger rewards (Silveira et al., 2020). The difference with our study might arise from the design of the experiment employed: in the Silveira et al. (Silveira et al., 2020) study, rats made decisions with two concurrent choices: stimulus duration of 1 second (lower reward) and 0.2 seconds (higher reward), while in our experiment, rats were presented with a single effort choice and had to decide whether to exert effort to categorize the stimulus to obtain the specific reward.

5-HTT^-/-^ exerted less cognitive effort at the beginning of discrimination but more at the end to reach similar discrimination accuracy compared with 5-HTT^+/+^ rats. This suggests a dynamic change in decisional anhedonia in rats lacking the 5-HTT. It could be that 5-HTT^-/-^ rats increased in learning rate during learning, as activation of serotonergic neurons increases the learning rate (Iigaya et al., 2018). 5-HTT^-/-^ might had a lower learning rate during stimuli discrimination at the beginning but may speed up the learning to catch up with 5-HTT^+/+^ rats later on. This later speed up learning might be in line with the finding that depressive patients make more accurate decisions to discriminate stimuli (Alloy and Abramson, 1979; Moore and Fresco, 2012). In sum, discrepancies in the relationship between the 5-HTT gene and depression (Ormel et al., 2019) might result from the interaction of increased anhedonia and learning rate.

The responsivity to the ambiguity of stimuli was subsequently measured during stimulus generalization. Both stimulus generalization and response time (RT) were analyzed during the stimulus generalization. 5-HTT^-/-^ and 5-HTT^+/+^ rats categorized the GSs and CSs similarly during decision-making under ambiguity. This indicates that the ablation of 5-HTT in rats didn’t change the capacity of generalization to process the ambiguity of the stimuli. The capacity of generalization between 5-HTT^+/+^ and 5-HTT^-/-^ mice was also similar in a touch screen-based task where the discrepancy of physical property (ambiguity) was the location of visual stimuli (Krakenberg et al., 2019a). These findings suggest that 5-HTT^-/-^ and 5-HTT^+/+^ rodents exert similar cognitive efforts to the categorization of the CSs and GSs during stimulus generalization. Notably, although the majority of stimulus generalization curves were monotonic graded responses, we observed 5 other types of generalization curves. Individual differences in stimulus generalization were also observed in humans (Stegmann et al., 2019). The individual differences in stimulus generalization might contribute to individual differences in cognitive efforts. Perhaps, a sophisticated computational model, rather than a linear regression model, can explain the different types of stimulus generalization curves.

During the decision-making before taking action, the response time (RT) plays a crucial role in information processing both in rats and humans (Donders, 1969; Blokland, 1998). Interestingly, we found that there was an interaction effect between genotype and session (learning) on RT to the stimuli. Overall, 5-HTT^+/+^ rats increased their RT with the progression of sessions, while 5-HTT^-/-^ rats tended to decrease the RT as the session increased. A human study proposes that the opportunity cost of time modulates cognitive effort: if the opportunity cost of time was high, subjects made more errors and responded faster (Otto and Daw, 2019). The observation that 5-HTT^-/-^ responded to the stimuli faster than 5-HTT^+/+^ rats might indicate that the opportunity cost of time was higher in 5-HTT^-/-^ rats. However, the response accuracy to the stimuli was similar between 5-HTT^-/-^ and 5-HTT^+/+^ rats. Applying computational models to analyze the data in a trial-by-trial manner might reveal whether the cognitive effort was different between 5-HTT^-/-^ and 5-HTT^+/+^ rats during stimulus generalization. Computational analysis in a human study revealed that pharmacological inhibition of 5-HTT reduced the cost of exerting effort (Meyniel et al., 2016). When processing reward information, depressed patients also responded to the stimuli faster (Goeleven et al., 2006). There could be a learning deficit in 5-HTT^-/-^ rats when processing generalization stimuli. As the generalization stimuli were not directly associated with rewards, 5-HTT^+/+^ rats responded to the stimuli progressive slower during the learning to reduce the opportunity cost for time and save energy (Boureau and Dayan, 2011). This learning effect, therefore, could affect decisional anhedonia, when the animals are learning to update the reward information to balance costs and benefits (Vessey, 1994; Der-Avakian and Markou, 2012).

Previous *in vivo* studies in rodents support the notion that serotonergic neurons encode the time for reward-related information processing. Serotonergic neurons have been found to fire tonically when the animal is waiting for the reward and then phasically when the animal received a reward (Li et al., 2016). Also, optogenetic activation of serotonergic neurons has been found to promote waiting (Fonseca et al., 2015). 5-HTT^-/-^ mice show under resting-state conditions a reduced firing rate of serotonergic neurons (Gobbi et al., 2001; Lira et al., 2003), which may lead to changes in waiting or the perception of reward. In our current study, the response time was different between 5-HTT^+/+^ rats and 5-HTT^-/-^ rats. 5-HTT^-/-^ rats responded to the generalization stimuli faster than 5-HTT^+/+^ rats as the session increased. These findings indicate that serotonin controls the time needed for reward-related information processing and represents an interesting substrate for the mechanistic understanding of decisional anhedonia before taking action.

As a limitation of the current experiment, we only included male rats. We, therefore, do not know whether our findings generalize to female 5-HTT^-/-^ rats. A pharmacological study showed that serotonin may play a minor role in cognitive effort in female rats (Silveira et al., 2020). Whether knocking out 5-HTT in female rats plays a role in cognitive efforts remains to be determined in the future.

In summary, 5-HTT ablation might increase decisional anhedonia during stimulus discrimination, and potentially reduce the cost of exerting cognitive effort to categorize generalization stimuli. This effect of 5-HTT ablation could be mild and interacts with learning. Therefore, the absence of the 5-HTT might contribute to the development of decisional anhedonia mildly. This partial effect could explain the discrepant findings of the link between the 5-HTT gene and depression.

## Acknowledgments

We thank Anthonieke Middelman for her assistance with animal breeding. The work is financially supported by a scholarship of the China Scholarship Council awarded to CG.

